# Sensory-modality independent activation of the brain network for language

**DOI:** 10.1101/714998

**Authors:** Sophie Arana, André Marquand, Annika Hultén, Peter Hagoort, Jan-Mathijs Schoffelen

## Abstract

The meaning of a sentence can be understood, whether presented in written or spoken form. Therefore it is highly probable that brain processes supporting language comprehension are at least partly independent of sensory modality. To identify where and when in the brain language processing is independent of sensory modality, we directly compared neuromagnetic brain signals of 200 human subjects (102 males) either reading or listening to sentences. We used multiset canonical correlation analysis to align individual subject data in a way that boosts those aspects of the signal that are common to all, allowing us to capture word-by-word signal variations, consistent across subjects and at a fine temporal scale. Quantifying this consistency in activation across both reading and listening tasks revealed a mostly left hemispheric cortical network. Areas showing consistent activity patterns include not only areas previously implicated in higher-level language processing, such as left prefrontal, superior & middle temporal areas and anterior temporal lobe, but also parts of the control-network as well as subcentral and more posterior temporal-parietal areas. Activity in this supramodal sentence processing network starts in temporal areas and rapidly spreads to the other regions involved. The findings do not only indicate the involvement of a large network of brain areas in supramodal language processing, but also indicate that the linguistic information contained in the unfolding sentences modulates brain activity in a word-specific manner across subjects.

## Introduction

Language can be realized in different modalities: amongst others through writing or speech. Depending on whether the sensory input modality is visual or auditory, different networks of brain areas are activated to derive meaning from the stimulus. Besides different brain circuits being recruited to process low-level sensory information, differences in linguistic features across sensory modalities prompt a differential activation of brain areas involved in higher-order processing as well. For instance, speech is enriched with meaningful prosodic cues, but also requires coarticulated signals to be parsed into individual words. Written text has the advantage of instantaneous availability of full information compared to the temporally unfolding nature of speech. These differences are paralleled in the brain’s response, and thus the sensory modality in which language stimuli are presented determines the dominant spatiotemporal patterns that will be elicited [19].

Regardless of low-level differences, the same core message can be conveyed in either modality. Therefore, language processing models of the past and present ([16], [18],[21]) include not only early sensory (up to 200ms) processing steps, but also contain late (200 - 500 ms), more abstract, and supposedly supramodal processing steps. While early processing is largely unimodal, and supported by brain regions in the respective primary and associative sensory areas, later processes (for instance lexical retrieval and integration) that activate several areas within the temporo-frontal language network are assumed to do so independent of modality.

In order to gain insight into the location and timing of brain processes representing this latter, higher order processing of the linguistic content, researchers so far relied on carefully manipulated experimental conditions. As a result, our current understanding of how the brain processes language across different modalities reflects a large variety in tasks (semantic decision task [10], error detection task [8], [11], passive hearing/listening [25], size judgment [31]) and stimulus material (words [10],sentences [3], and stories [5, 14, 38]). Despite this wealth of experimental findings and resulting insights, an important interpretational limitation stems from the fact that the majority of studies employ modality specific low-level baseline conditions (tone pairs and lines, spectrally rotated speech and false fonts, non-words, white noise [29]) to remove the sensory component of the processing. It is difficult to assess in how far such baselines are comparable across auditory and visual experiments. Recent fMRI work has demonstrated sensory-modality independent brain activity by directly comparing the BOLD response across visual and auditory presentations [14, 38]. Yet, fMRI signals lack the temporal resolution to allow for a temporally sufficiently fine-grained investigation of the response to individual words.

Few studies used magnetoencephalography (MEG) to study supramodal brain activity and all are based on event-related averaging [31, 3, 36, 47]. Averaged measures capture only generic components in the neural response. While generic components make a large contribution to the neural activity measured during language processing, there also exist meaningful variability in the neural response that is stimulus-specific and robust [4]. A complete analysis of the supramodal language network needs to tap into these subtle variations as well.

Here, we overcome previous limitation by achieving a direct comparison without relying on modality-specific baseline conditions, leveraging word-by-word variation in the brain response. Using MEG signals from 200 subjects, we performed a quantitative assessment of the sensory modality independent brain activity following word onset during sentence processing. The MEG data forms part of a large publicly available dataset [40], and has been used in other publications [28], [39], [24], [27]. We identified widespread left hemispheric involvement, starting from 325 ms after word onset in the temporal lobe and rapidly spreading to anterior areas. These findings provide a quantitative confirmation of earlier findings in a large study sample. Importantly, they also indicate that supramodal linguistic information conveyed by the individual words in sentence context leads to subtle fluctuations in brain activation patterns that are correlated across different subjects.

## Materials and Methods

### Subjects

A total of 204 native Dutch speakers (102 males), with an age range of 18–33 years (mean of 22 years), participated in the experiment. In the current analysis, data from 200 subjects were included. Exclusion of four subjects was due to technical issues during acquisition, which made their datasets not suitable for our analysis pipeline. All subjects were right-handed, had normal or corrected-to-normal vision, and reported no history of neurological, developmental, or language deficits. The study was approved by the local ethics committee (CMO, the local “Committee on Research Involving Human Participants” in the Arnhem–Nijmegen region) and followed the guidelines of the Helsinki declaration. All subjects gave written informed consent before participation and received monetary compensation for their participation.

### Experimental Design

The subjects were seated comfortably in a magnetically shielded room and presented with Dutch sentences. From the total stimulus set of 360 sentences six subsets of 120 sentences were created. This resulted in six different groups of subjects who were presented with the same subset of stimuli although in a different (randomized) order with some overlap of items between groups. Within each group of subjects half of them performed the task in only the visual, the other half in only the auditory modality. In the visual modality, words were presented sequentially on a back-projection screen, placed in front of them (vertical refresh rate of 60 Hz) at the center of the screen within a visual angle of 4 degrees, in a black mono-spaced font, on a grey background. Each word was separated by an empty screen for 300 ms and the inter-sentence interval was jittered between 3200 and 4200 ms. Mean duration of words was 351 ms (minimum 300 ms and maximum 1400 ms), depending on word length. The median duration of whole sentences was 8.3 s (range 6.2 - 12 s). Auditory sentences had a median duration of 4.2 s (range 2.8 - 6.0 s) spoken in a natural pace. The duration of each visual word was determined by the following quantities: (i) the total duration of the audio-version of the sentence/word list (audiodur), (ii) the number of words in the sentence (nwords), (iii) the number of letters per word (nletters), and (iv) the total number of letters in the sentence (sumnletters). Specifically, the duration (in ms) of a single word was defined as: (nletters/ sumnletters) * (audiodur + 2000–150 * nwords). In the auditory task the stimuli were presented via plastic tubes and ear pieces to both ears. Before the experiment, the hearing threshold was determined individually and the stimuli were then presented at an intensity of 50 dB above the hearing threshold. A female native Dutch speaker recorded the auditory versions of the stimuli. The audio files were recorded in stereo at 44100 Hz. During the post processing the audio files were low-pass filtered at 8500 Hz and normalized so that all audio files had the same peak amplitude, and same peak intensity. All stimuli were presented using the Presentation software (Version 16.0, Neurobehavioral Systems, Inc). Sentences were presented in small blocks, of five sentences each, along with blocks containing scrambled sentences, which were not used here. See Lam et al. [28] for more details about the stimulus material used. In order to check for compliance, 20% of the trials were randomly followed by a yes/no question about the content of the previous sentence/word list. Half of the questions addressed the content of the sentence (e.g. Did grandma give a cookie to the girl?) whereas the other half, addressed one of the main content words (e.g. Was the word ‘grandma’ mentioned?). Subjects answered the question by pressing a button for ‘Yes’/ ‘No’ with their left index and middle finger, respectively.

### MEG Data Acquisition & Structural imaging

MEG data were collected with a 275 axial gradiometer system (CTF). The signals were analog low-pass-filtered at 300 Hz and digitized at a sampling frequency of 1,200 Hz. The subject’s head was registered to the MEG-sensor array using three coils attached to the subject’s head (nasion, and left and right ear canals). Throughout the measurement, the head position was continuously monitored using custom software [44]. During breaks the subject was allowed to reposition to the original position if needed. Participants were able to maintain a head position within 5 mm of their original position. Three bipolar Ag/AgCl electrode pairs were used to measure the horizontal and vertical electrooculogram and the electrocardiogram.

A T1-weighted magnetization-prepared rapid gradient-echo (MP-RAGE) pulse sequence was used for the structural images, with the following parameters: volume TR = 2300 ms, TE = 3.03 ms, 8 degree flip-angle, 1 slab, slice-matrix size = 256 × 256, slice thickness = 1 mm, field of view = 256 mm, isotropic voxel-size = 1.0 × 1.0 × 1.0 mm. A vitamin-E capsule was placed as fiducial behind the right ear to allow a visual identification of left-right consistency.

### Preprocessing

Data were bandpass filtered between 0.5 and 20 Hz, and epoched according to sentence onset, each epoch varying in length, depending on the number of words within each sentence. Samples contaminated by artifacts due to eye movements, muscular activity, and superconducting quantum interference device jumps were replaced by NaN before further analysis. Since all sentences had been presented in random order, we reordered sentences for each subject to yield the same order across subjects. Subsequently, the signals of the auditory subjects were temporally aligned to the signals of the visual subjects, ensuring coincidence of the onset of the individual words across modalities (Figure 1A). This alignment was needed to accommodate for differences in word presentation rate. The alignment was achieved by first epoching the auditory subject’s signals into smaller overlapping segments. Each segment’s first sample corresponded to one of the word onsets as annotated manually according to the audio file while each segment’s length depended on the duration of the visual presentation of the corresponding word. Finally, all segments were concatenated again in the original order. By defining segments that were longer than the corresponding auditory word duration, the neural response to each word is fully taken into account and matched to the visual signal, even in the case of short words where the response partly coincided with the next word presentation. MEG data were then downsampled to 120 Hz.

**Figure 1.**
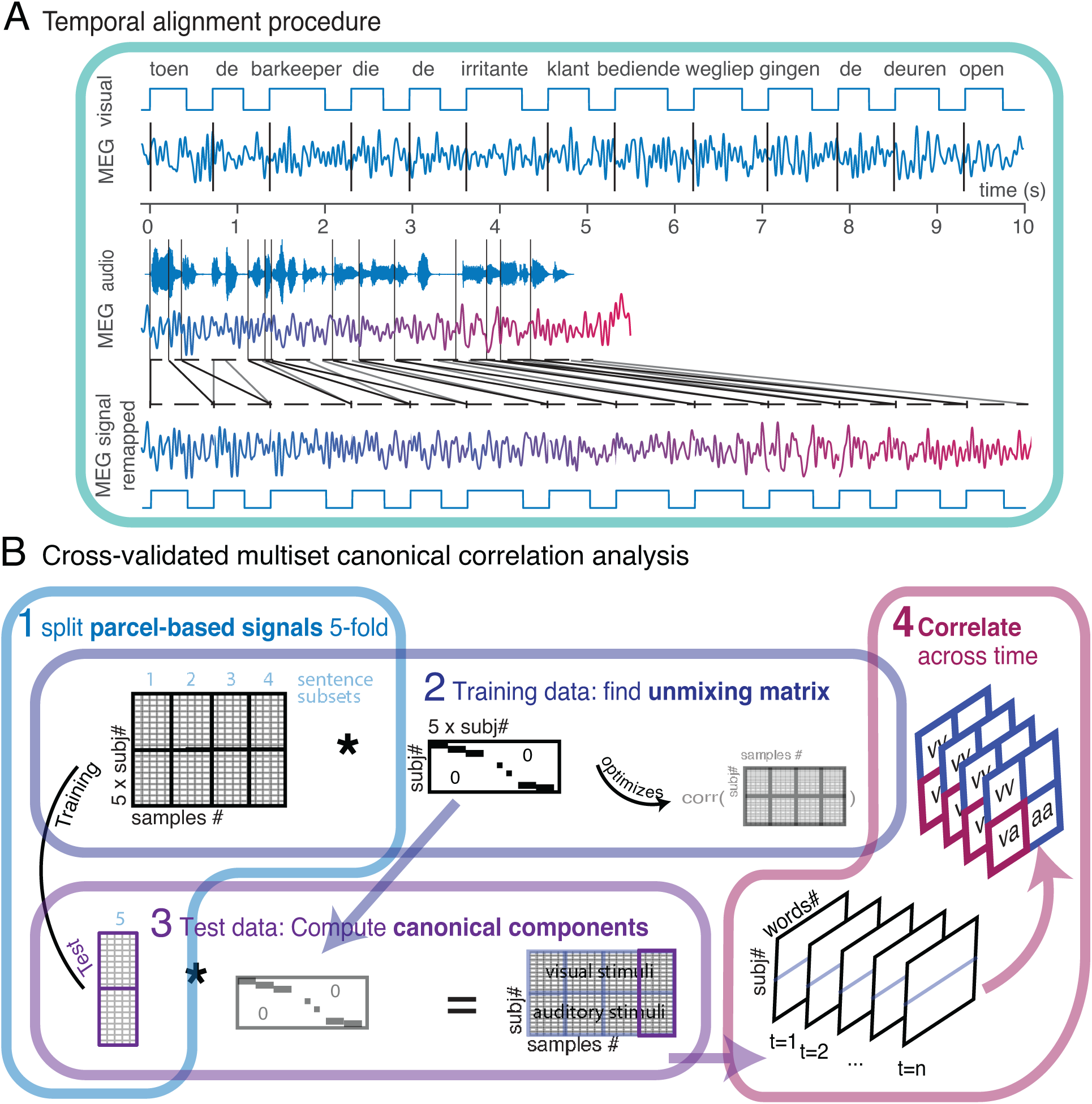
Analysis pipeline. (A) Temporal alignment procedure. MEG signals of auditory and visual subjects differed in length due to different presentation rates. To achieve alignment between signals of auditory and visual subjects, auditory signals were epoched into overlapping segments. Each segment’s first sample corresponds to the auditory word onset but each segment’s length depends on the duration of the equivalent visual stimulus. Segments were then concatenated in original order to recover signal for the full sentence length. This way, the neural response to each word is fully taken into account in further comparisons, including in the case of short words for which stimulus late processing coincided with the next word presentation. (B) Starting point for the multi-set canonical correlation analysis were parcel-based neural signals for all subjects, consisting of five spatial components each. 1. Signals for all sentence trials were split into five subsets and for cross-validation one subset of sentences was left out as test data, while the remaining four subsets served as training data. 2. Based on the training dataset only an unmixing matrix was found, per parcel, defining the linear combination of the five spatial components so that the correlation across sets (subjects) and time samples were maximized. The cross-covariance was computed between all subjects’ spatial components and across time collapsing over sentence trials. 3. The projection was applied to the test data to compute canonical variables for the left out sentence trials (purple outline) for all subjects. Steps 2 and 3 were repeated for all folds until each sentence subset had been left out once and the resulting canonical variables were concatenated until the entire signal was transformed. 4. Canonical variables were epoched according to word onsets and for each time point a subject-by-subject correlation matrix was computed across words. Correlation between cross-modal subjects (pink outline) were interpreted as quantifying supramodal activation.

### Source Reconstruction

We used linearly constrained minimum variance beamforming (LCMV) [46] to reconstruct activity onto a parcellated cortically constrained source model. For this, we computed the covariance matrix between all MEG-sensor pairs, as the average covariance matrix across the cleaned single trial covariance estimates. This covariance matrix was used in combination with the forward model, defined on a set of 8,196 locations on the subject-specific reconstruction of the cortical sheet to generate a set of spatial filters, one filter per dipole location. Individual cortical sheets were generated with the Freesurfer package [12, version 5.1] (surfer.nmr.mgh.harvard.edu), coregistered to a template with a surface-based coregistration approach, using Caret software [45] (download here and here), and subsequently downsampled to 8,196 nodes, using the MNE software [17] (martinos.org/mne/stable/index.html)). The forward model was computed using FieldTrip’s singleshell method [34], where the required brain/skull boundary was obtained from the subject-specific T1-weighted anatomical images. We further reduced the dimensionality of the data to 191 parcels per hemisphere [39]. For each parcel, we obtained a parcel-specific spatial filter as follows: We concatenated the spatial filters of the dipoles comprising the parcel, and used the concatenated spatial filter to obtain a set of time courses of the reconstructed signal at each parcel. Next, we performed a principal component analysis and selected for each parcel the first five spatial components explaining most of the variance in the signal.

### Multi-set Canonical Correlation Analysis

Multi-set canonical correlation analysis (MCCA) [37], [13] was applied to find projections of those five spatial components that would transform the subject-specific signals so as to boost similarities between them. Canonical correlation analysis (CCA) is a standard multivariate statistical method often used to investigate underlying relationships between two sets of variables. Classically, canonical variates are estimated by transforming the two sets in a way that optimizes their correlation. We applied a generalized version of the classical approach (MCCA) [26], which extends the method to multiple sets, here multiple subjects. In our case, we find linear combinations of the five spatial components for each of two subjects, so that the correlation across time between those subjects is maximized. Since we have more than two subjects, we find for each subject its own linear combination, which maximizes the correlation across time between all subjects from both modality groups (auditory and visual stimulation). Following Parra we obtained the optimal projection as the eigenvector with the largest eigenvalue of a square matrix D^-1^R, where R and D are square matrices:

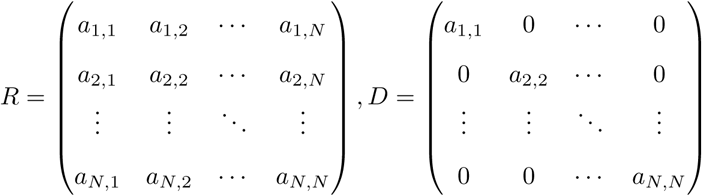

Where a^lk^ are cross-covariance matrices between subject pairs and D contains only the diagonal blocks of within subject covariances [37]. In our case the cross-covariance matrices are of size 5-by-5 containing the cross-covariance between all five spatial components for a given subject (pair). The cross-correlation is computed across time points for each sentence and subsequently averaged across sentences. It is important to note here, that the canonical variates resulting from the optimal projection do not reflect sentence averages anymore but have the same temporal resolution as the original source signals. CCA is prone to overfitting and known to be unstable [15]. For reliable CCA estimates the number of samples should be much larger than the number of features, i.a. a sample-to-feature ratio of 20/1 is recommended [43]. We estimated the canonical variables over concatenated data, which included between 756 and 1453 samples per sentence compared to only five features (spatial components) which provides a decent sample-to-feature ratio. Further, we estimated our canonical variables out-of sample using 5-fold cross-validation to limit overfitting. We randomly split all sentences into five subsets, estimating projections on 96 sentences and applying them to the 24 left out sentences (Figure 1B).

### Statistical Analysis

As per the study design, the subjects were assigned to one of six stimulus sets. Different groups of subjects were presented with different sets of sentences. Since MCCA relies on commonalities across datasets, we could only combine data from subjects who received the exact same stimulation. We therefore applied MCCA for each subgroup of subjects who listened to or saw the same stimuli separately. Initially, we constrained our analysis to the first set of 33 subjects (henceforth exploratory dataset). After applying the projection to the data we computed a time-resolved Pearson correlation between all possible subject pairings. To this end, we first epoched the resulting canonical components according to individual word onsets and selected only content words (nouns, adjectives & verbs) for subsequent steps. Before computing the correlation we subtracted the mean across samples. For each pair of subjects, we computed the correlation between two sets of observations, i.e. a pair of vectors with each data point reflecting the subject-specific neural signal for each of the individual words (lexical items), at a given time point relative to word onset, and at a given cortical location. Correlation coefficients of cross-modality pairings, that is correlations between subjects reading and subjects listening to the sentences are interpreted as capturing supramodal processing. We used a permutation test with clustering over time and space (parcels) for family-wise error rate correction for statistical inference (Maris and Oostenveld, 2007), using 1000 randomizations of the epoched words. To this end, we randomized word order for the source-reconstructed parcel time series of the auditory subjects to test for exchangeability of the exact word pairing across sensory modalities. By destroying the one-to-one mapping of individual lexical items, the null distribution allowed for a distinction between individual item specific shared variance, and shared variance due to a more generic response. For a more conservative test, we additionally tested the observed correlation patterns against a null distribution obtained by permuting sentence order and doing this prior to MCCA. The results and a description of this additional test can be found in supporting Figure 3-1.

We then analyzed the remaining sets of subjects (confirmatory dataset) using the same analysis pipeline. We evaluated the overlap in the results across all six subgroups using information prevalence inference [1]. Prevalence inference allows to formulate a complex null hypothesis, i.e. that the prevalence of the effect is smaller than or equal to a threshold prevalence, where the threshold can be realized by different values. For each of the six sets of data, we obtained spatial maps of time-resolved supramodal correlations, as well as 1000 permutation estimates after word order shuffling (see above). We used the smallest observed average correlation across subgroups as the second-level test statistic. We then tested the majority null hypothesis of the prevalence of the effect being smaller or equal to a threshold prevalence. For this, we computed the largest threshold such that the corresponding null hypothesis could still be rejected at the given significance level *α*. This was done after concatenating the minimum statistic from all parcels and time points, using the maximum statistic to correct for multiple comparisons in time and space (parcels). For each parcel we evaluated the highest threshold at which the prevalence null hypothesis could be rejected at a level of *α* = 0.05 (see supporting Figures 4-1 and 4-2 for cortical maps showing thresholds averaged and per time point).

### Code Accessibility

All analyses were done with custom-written MATLAB scripts and FieldTrip [35] and the corresponding code is available upon request.

## Results

### Modality-specific activation

We first quantified the similarity between different subjects’ brain response within only the exploratory dataset (33 subjects) by correlating word-by-word fluctuations in brain activity between all possible pairs of subjects. Averaging the correlations across those subject pairings for which subjects were stimulated either in the same sensory modality, or each in a different modality, allowed us to evaluate the modality-specific brain response and the supramodal response, respectively. As displayed in Figure 2, early sensory cortical areas only show correlated activity for the group of subjects receiving the stimuli in the corresponding sensory modality, for the visual (red), and auditory (blue) modalities. We found that MCCA is a crucial analysis step in order to reveal meaningful inter-subject correlations. Only after MCCA does cortical activity in visual and auditory areas become significantly correlated across those subjects performing the task in the visual or auditory domain respectively (Figure 2A).

**Figure 2.**
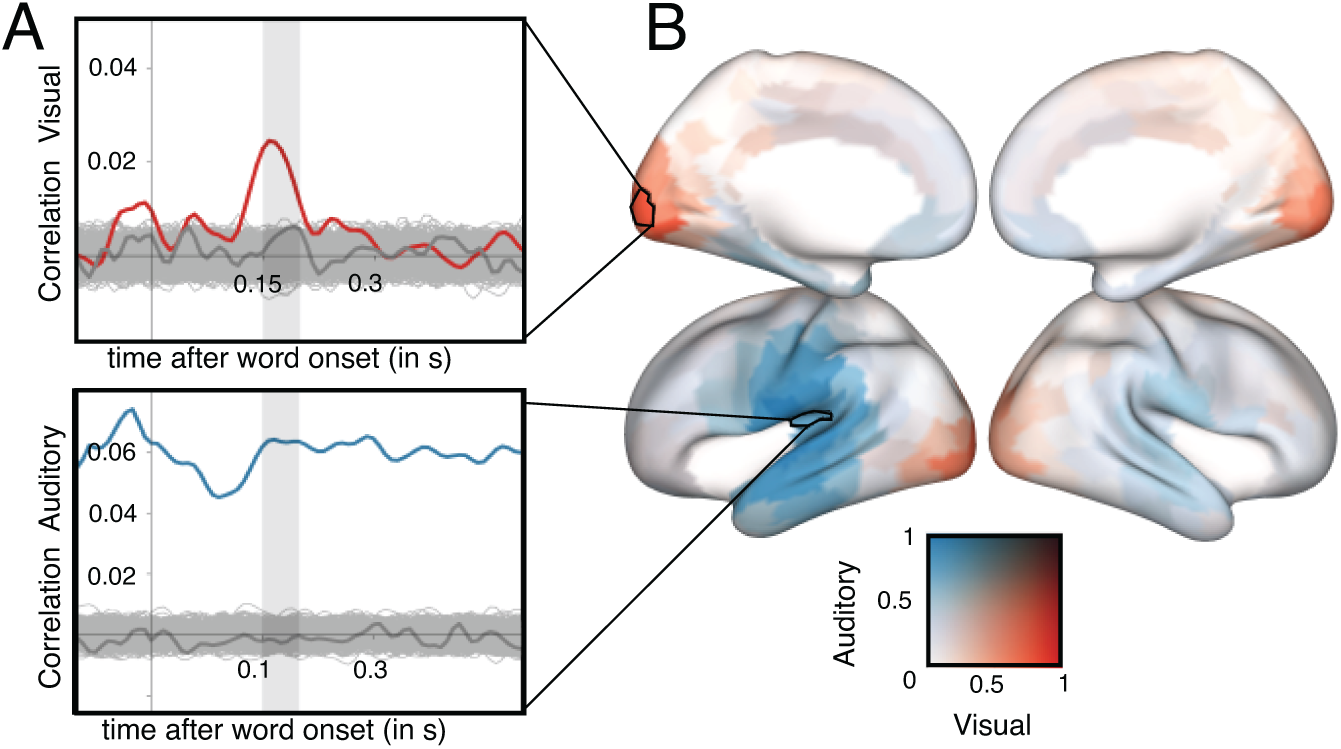
Specificity of the within-modality correlated activity patterns. (A) Time-resolved correlation values averaged across all visual subject pairings for a parcel in left primary visual cortex (upper panel) and all auditory subject pairings for a parcel in left primary auditory cortex (lower panel), before (dark grey line), and after MCCA (blue and red lines). Light grey lines show recomputed correlation values for 1000 random permutations of word order across subjects. Notably, signals of auditory subjects highly correlate even before word onset. This is likely due to a more varied distribution of information in the auditory signal caused by the continuous nature of auditory stimulation and as a result differing time points at which individual words become uniquely recognizable. The MCCA is blind to the stimulus timing and will thus find canonical variables that yield maximal correlations at any timepoint if possible. (B) Cortical map of the spatial distribution of correlations, comparing within modality visual subject pairs (red) with auditory subject pairs (blue). Correlation strength is expressed as the Pearson correlation coefficient averaged over a time window from 150 to 200 ms post word-onset and normalized by the maximum value of that window.

### Supramodal activation patterns

We averaged between-subject correlations over all cross-modal subject pairings as a metric for supramodal activity. We observed significant supramodal correlated activation patterns in mostly left-lateralized cortical areas (Figure 3). The effect has a large spatial and temporal extent, becoming apparent as early as 250 ms and lasting until 700 ms after word onset. Parcels in middle superior temporal gyrus (STG) contribute to the effect at the earliest time points, followed by the posterior and anterior part of the STG and about 50 ms later the anterior temporal pole. Supramodal correlated activation in ventral temporal cortex follows a similar temporal and spatial pattern, with supramodal correlations starting out more posterior around 292 ms and evolution towards the middle anterior temporal lobe at 308 ms. Other areas that express supramodal activity at relatively early time points are medial prefrontal cortex and primary auditory cortex (250 ms), followed by subcentral parietal regions and supramarginal gyrus at around 300 ms, and finally dorsolateral frontal cortex (DLFC, 325 ms). By the time 375 ms have passed, the entire orbito-frontal cortex, anterior and DLFC as well as inferior frontal gyrus (IFG) show strong supramodal subject correlation. Supramodal activation in the frontal lobe further extends towards posterior regions including pre- and postcentral gyrus. At around 400 ms supramodal subject correlation in the anterior temporal pole reaches its peak. In addition to the lateral cortical areas, correlated activity also extends to left dorsal and ventral anterior cingulate cortex (ACC) as well as left fusiform gyrus. The spatio-temporal patterns of supramodal activation described so far are robust, also when ordinal word position and MCCA overfitting is controlled for in the statistical evaluation (see supporting Figure 3-1).

**Figure 3.**
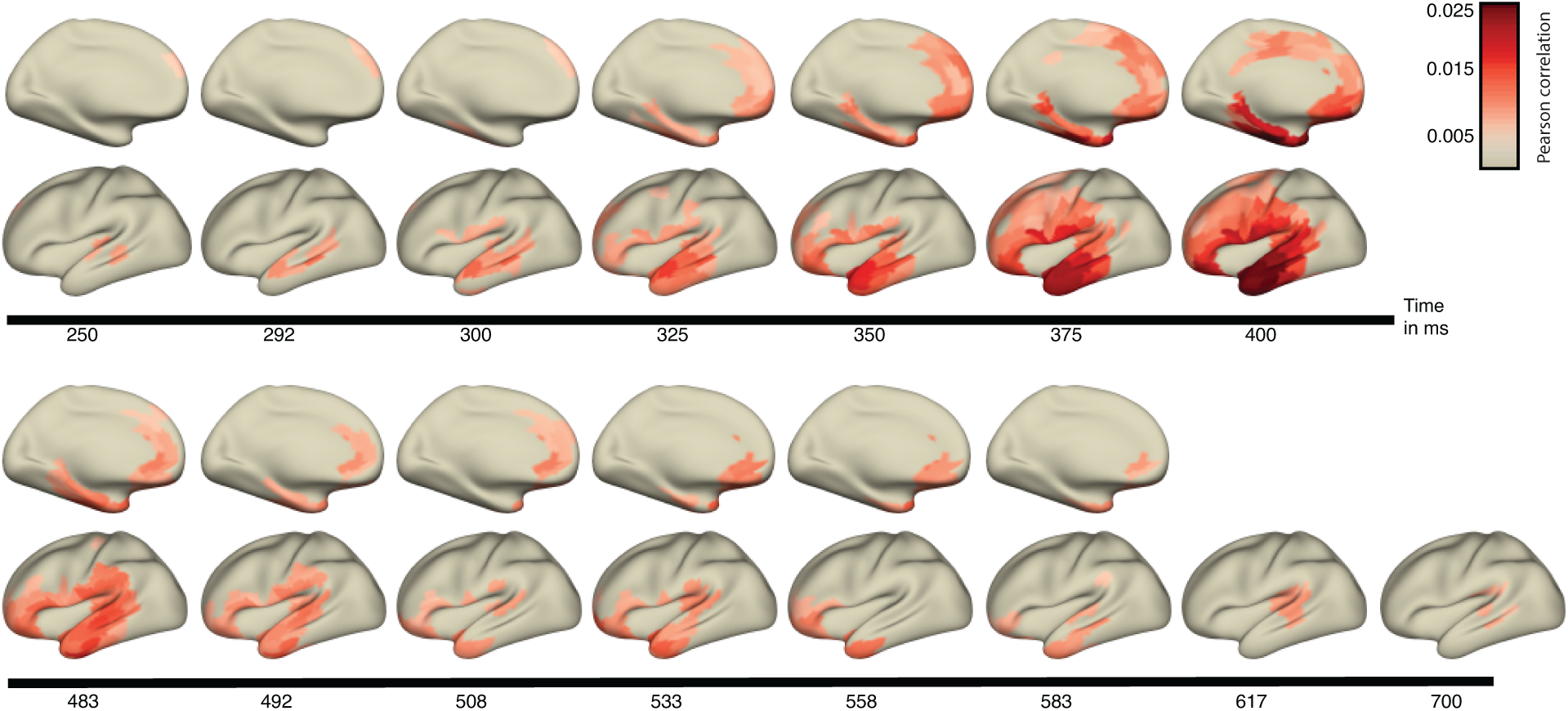
Supramodal correlated activity patterns. Time-resolved spatial maps of supramodal correlated activity patterns (averaged over all possible cross-modal subject pairings) in the left hemisphere. Medial views of the brain surface are depicted in the first and third row, lateral views in the second and fourth row. Color codes for strength of correlation. Colored parcels were most strongly correlated between cross-modal subject pairs (nonparametric permutation test, corrected for multiple comparisons).

### Prevalence Inference

Our confirmatory analysis combined over all six datasets and tested whether the spatiotemporal patterns observed in the exploratory dataset would generalize to the population. For those parcels at which the global null hypothesis could be rejected, we infer that at least in one of the datasets an effect of supramodal processing was present (Figure 4). In addition, we evaluated the majority null hypothesis of whether, in the majority of subgroups in the population, the data contains an effect (threshold > 0.5, significant parcels under the majority null hypothesis outlined in black in figure 4B).

**Figure 4.**
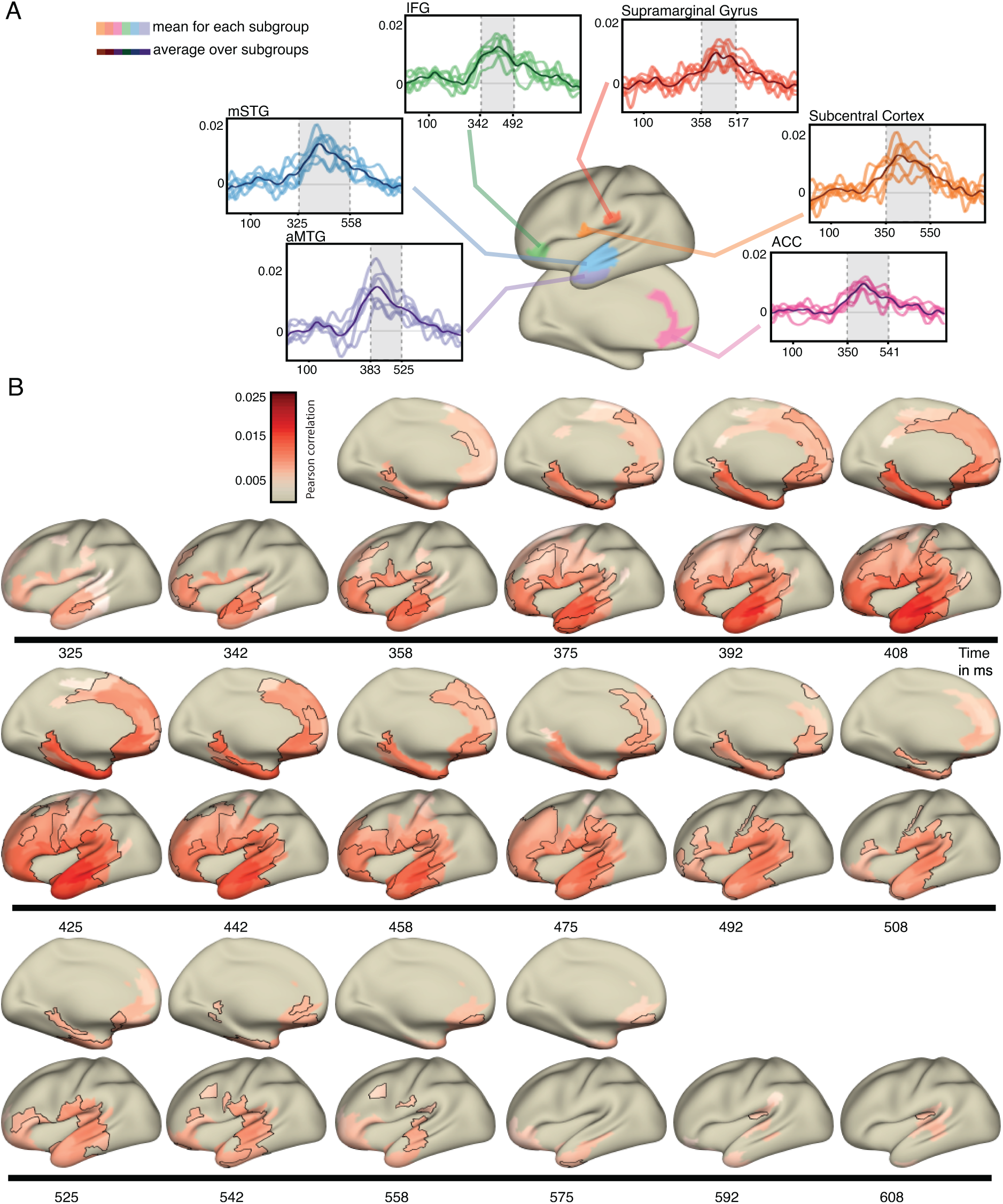
Supramodal correlated activity patterns consistent across the majority of datasets. Supramodal correlated activity patterns of word-specific activity consistent across the majority of datasets. (A) Averaged correlation time courses (mean over all possible cross-modal subject pairings) are shown for selected parcels in inferior frontal Gyrus (IFG, green), supramarginal Gyrus (red), Subcentral Cortex (orange), Anterior cingulate cortex (ACC, pink), anterior middle temporal Gyrus (aMTG, purple), and middle superior temporal gyrus (mSTG, blue). Time courses are shown for each dataset individually (light-colored lines) as well as averaged (dark lines). Grey shaded areas mark statistically significant time points. (B) Time-resolved spatial maps of cross-modal correlations in the left hemisphere. Medial views of the brain surface are depicted in the first, third and fifth row, lateral views in the second, fourth and sixth row. For those parcels that were part of the largest nominal suprathreshold cluster tested on only the exploratory dataset, the mean correlation over all six datasets is shown. Color codes for strength of correlation. In addition, the parcels at which the majority null hypothesis according to prevalence inference could be rejected are outlined in black.

The global null hypothesis (no information in any set of subjects in the population) could be rejected at a level of *α* = 0.05 in on average 40 parcels per time point (between 325 and 617 ms after onset, std. = 31.48). For those parcels, the average largest lower bound *γ*0 at which the prevalence null hypothesis can be rejected is shown in supporting Figures 4-1 and 4-2. For those parcels for which the largest bound *γ*0 is larger than or equal to 0.5 we can infer that in the majority of the datasets the activity patterns were similar across subjects independent of modality.

This majority null hypothesis could be rejected (at a level of *α* = 0.05) in 90% of parcels that also showed a global effect (Figure 4, see supporting Figure 4-3 for results in the right medial hemisphere). Compared to the temporal pattern of the largest nominal suprathreshold cluster from the cluster-based permutation test conducted on the exploratory dataset, the effect became significant in the majority of datasets later in time and was less long lasting (325 - 608 ms). Given this time span, the majority null hypothesis was rejected in on average 42% of those parcels contributing most to the largest cluster. The orbitofrontal cortex and IFG showed an involvement in supramodal processing in both analyses, but the effect there was much more temporally sustained in the exploratory dataset. In addition, according to the exploratory dataset, supramodal activation of STG occurred almost 100 ms earlier as compared to IFG. Based on the confirmatory dataset, however, supramodal correlated activation in IFG and STG appeared almost simultaneously. Finally, the exploratory analysis revealed supramodal activation in primary and premotor areas extending over the entire left dorsolateral surface, of which only the most ventral parcels close to the Sylvian fissure were significantly supramodal in the majority of datasets. Thus, the spatial extent of the effect was partly reduced for prevalence inference, compared to the cluster-based permutation approach on the exploratory data. Nevertheless, widely overlapping anatomical regions were indicated by both analyses, encompassing dorsolateral frontal gyrus and the middle & superior parts of the temporal lobe at first and later also inferior frontal and orbito-frontal cortex, as well as anterior temporal lobe.

## Discussion

Our aim was to quantify similarities of the brain response across reading and listening at a fine temporal scale. To this end we correlated word-by-word fluctuations in the neural activity across subjects receiving either auditory or visual stimulation. We identified a widespread left-lateralized brain network, activated independently of modality starting 325 ms after word onset. Importantly, dividing our large study sample into six subsets, we could directly quantify the consistency and generalizability of these activity patterns. The spatial distribution of the supramodal activation is in line with the known involvement of left hemispheric areas, including parts of left temporal cortex, left inferior parietal lobe, as well as prefrontal cortex [48], [10], [11], [7], [30], [29], [42], [23]. The involvement of both STG and IFG fits predictions from the Memory, Unification and Control model (MUC), in which activity reverberating within a posterior-frontal network ([2], [18]) is thought to be crucial for language processing. According to the MUC model temporal and parietal areas support the retrieval of lexical information, while unification processes are supported by inferior frontal cortex. Bidirectional communication [39] between these areas is facilitated by white matter connections. We observe that temporal areas are supramodally activated at earliest time points and sustain activation for the longest compared to other regions. Over time, supramodal activation spreads from middle and posterior left STG to the anterior temporal pole. This rapid progression of activity from posterior to anterior regions mirrors previous observations ([31], [47]), adding to those findings a direct quantitative comparison of the supramodal brain activity.

### Beyond the core language network and the single word level

We observed modality-independent activity in dorsal frontal cortex, in addition to more widely reported inferior parts of the frontal cortex [25], [29], [11], [31], [23], [33]. This could be due to us using linguistically rich sentence material, of varying syntactic complexity, as opposed to single words [10],[31],[6],[30],[47] or short phrases [3],[8],[7]. Indeed, discrepancies with respect to frontal lobe involvement in modality-independent processing seem to mainly arise from differences in stimulus material and task demands [7]. A recent meta-analysis has identified that more complex syntax robustly activates dorsal parts of the left IFG [20]. Further, a previously published analysis of these MEG data showed DLFC to be sensitive to sentence progression effects [24]. Two previous fMRI studies using narratives [14, 38] add to the debate. Regev et al. correlated BOLD responses evoked by different modalities. They report supramodal activation in the left frontal lobe, extending beyond inferior frontal regions. Deniz et al. study modality-independent brain areas by modelling semantic features of the stimulus in one modality and used the model to predict the BOLD signal in the other modality. They report BOLD signals in prefrontal cortex to be well predicted across modalities. In summary, while complex stimuli consistently activate prefrontal areas beyond inferior frontal cortex, the exact stimulus features which cause this supramodal activation are still debated.

Some previous studies, using narratives and fMRI, report supramodal activation not to be restricted to the left hemisphere [14, 38, 25]. It could be that the previously observed bilateral involvement is due to differences in context-based semantic processing during narratives, as compared to the processing of isolated sentences in our experiment. Menteni and colleagues have specifically contrasted BOLD activity in response to sentences presented within a neutral or a local context. The authors indeed reported right frontal cortex to be more sensitive to local discourse context as compared to its left-hemispheric homolog [32]. Further research is needed to determine whether this effect of presence and absence of narrative thread similarly affects lateralisation of brain activity in MEG.

Even though mostly restricted to the left hemisphere, our results also implicate extra-linguistic areas in supramodal processing. Specifically, we find bilateral supramodal activation within ACC. The ACC is a midline structure, forming part of a domain-general executive control network supporting language processing [18], [9]. It is sensitive to statistical contingencies in the language input and thus might play a role in mediating learning and adaptation in response to predictive regularities in both local experimental as well as global environment [49]. It should be noted that deep sources are normally poorly detectable in MEG [22] and we thus consider any interpretations with respect to the midline structures as tentative.

### Supramodal orthography-phonology mapping

We observed supramodal activation in post-central and subcentral gyrus, as well as supramarginal gyrus, which coincides temporally with supramodal activation of primary auditory cortex. Activity in supramarginal gyrus has been repeatedly elicited by cross-modal tasks [41], such as rhyming judgments to visually presented words [6], for which conversion between orthographic and phonological representations is likely needed. At the same time, post- and subcentral areas partly span articulatory motor and somatosensory areas for the mouth and tongue. Together, the supramodal activation of these areas suggests that retrieval of phonetic and articulatory mappings is not limited to speech perception only but also occurs during passive reading.

### Leveraging word-by-word variability of the neural response

Neuroelectric brain signals exhibit strong moment-to-moment variability. While some of this variability is related to the experimental stimulation, and therefore associated with specific cognitive activity, some of it is unrelated, ongoing neural activity. By applying MCCA across subjects we reduced this type of noise and made subtle word-by-word fluctuations in the MEG signal interpretable. Comparing neural activity across subjects is challenging due to differing position or orientation of neuronal sources relative to the MEG sensors. We used parcellated MEG source reconstruction in combination with exact temporal alignment of individual sentences across subjects. This allowed for the extraction of signal components that are shared across subjects, thus reducing the intersubject spatial variability, which is commonly observed in more traditional (for instance, dipole fitting) procedures [47]. MCCA thus allowed us to more directly investigate time-resolved inter-subject correlations and move beyond event-related averages [31]. Importantly, our analysis approach allows us to conclude that the identified supramodal activity is word-specific. Our findings therefore go beyond showing a general activation of those areas as compared to baseline but rather reveal consistent word-by-word fluctuations of activation within the recruited areas.

### Latency of supramodal processing

The temporal alignment procedure, as a necessary preparation step for the MCCA procedure, followed by the estimation of time-resolved intersubject correlations, focused on common signal aspects that are exactly synchronized across subjects. The differences in sensory modality specific characteristics of the input signal require dedicated processing with likely different processing latencies, which may also lead to latency differences in the activation of supramodal areas. For example, Marinkovic and colleagues report shorter reaction times during the visual task, yet found earlier activity peaks for the auditory task in corresponding early sensory cortex and left anterior temporal lobe [31]. In contrast, other work observed earlier anterior temporal lobe activation for visual, compared to auditory stimulation [3]. Our results indicate a certain degree of overlap across modalities in the temporal window within which supramodal cortical areas are activated. It is possible, that we observed more temporally extensive activation, for instance, related to unification processes, because we used longer sentences. In addition, any overlap may have been amplified as a necessary consequence of the MCCA procedure. Evidently, correlations between signals from auditory subjects were boosted with less temporal specificity compared to visual subjects (Figure 2B). This observation was unexpected and may be due to more continuous stimulation in the auditory experiment. As the sound of a spoken word unfolds, the timing at which it becomes uniquely recognizable will vary across word. Thus, the distribution of information in the auditory signal is much more varied as compared to the visual. MCCA will pick up on any common relationship across subjects regardless of timing. In our specific application, projections were estimated on concatenated data, effectively making the method blind to word onset boundaries.

In conclusion, this study provides direct neurophysiological evidence for sensory modality independent processes supporting language comprehension in multiple left hemispheric brain areas. We identified a network of areas including domain general control areas as well as phonological mapping circuits over and above traditional higher-level language areas in frontal and temporal-parietal regions, by quantifying between-subject consistency of their respective word-specific activation patterns. These consistent activation patterns were word-specific, and thus likely reflect more than just generic activation during language processing. Finally, we show that alignment of individual subject data through MCCA is a promising tool for investigating subtle word-in-context specific modulations of brain activity in the language system.

## Acknowledgments

We thank Phillip Alday for providing helpful comments. This work was supported by The Netherlands Organisation for Scientific Research (NWO Vidi: 864.14.011, awarded to J.M.S.).

---

**Figure 3-1.**
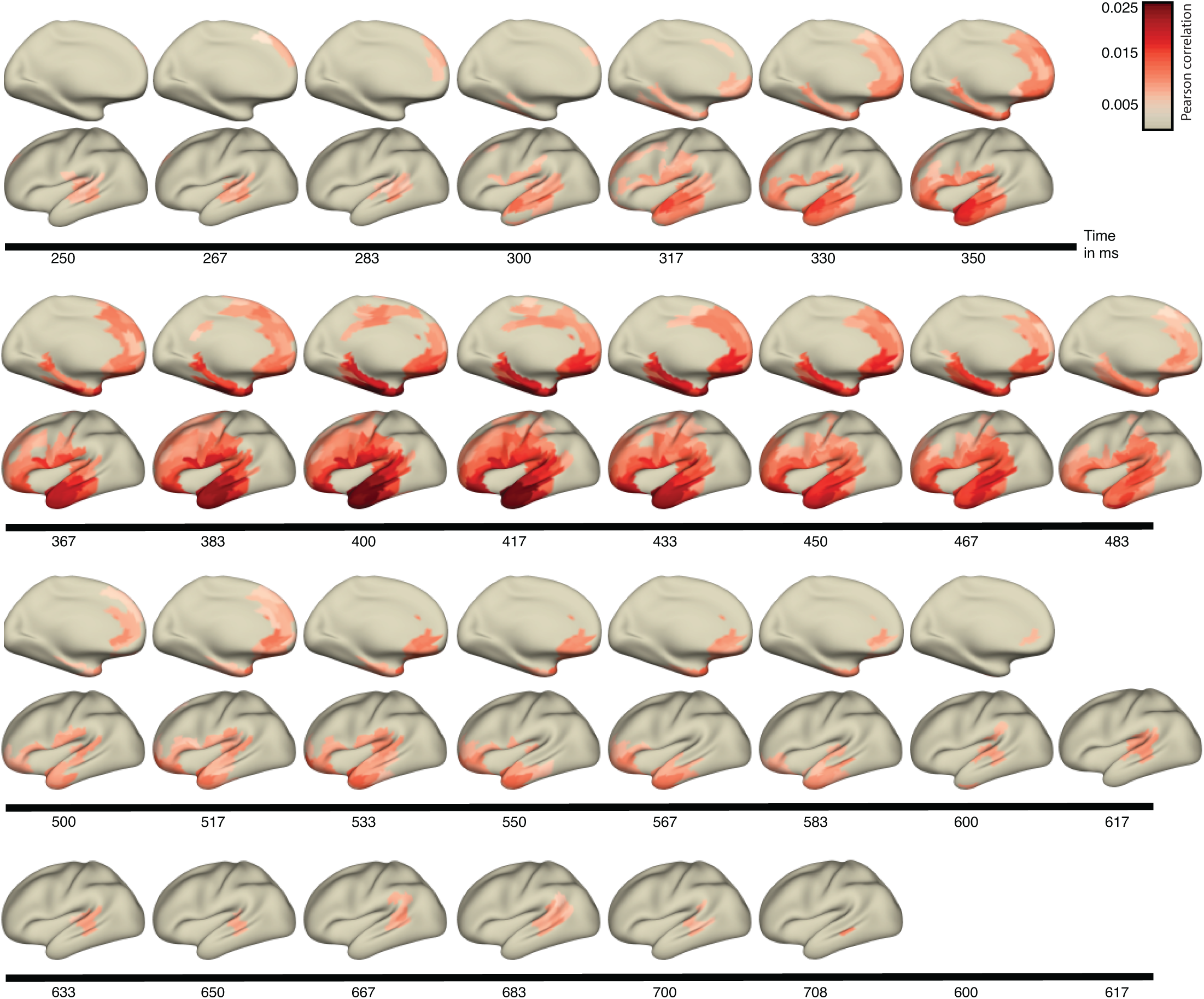
Figure 3-1. Significant supramodal correlated activity patterns as assessed by additional permutation test. In an additional significance test, we shuffled the sentence order 500 times prior to MCCA, controlling for the possibility that MCCA as a preprocessing step may artificially increase correlations between subjects (through overfitting). Importantly, this permutation was not fully unconstrained, since we aimed at aligning sentences across modalities with the same number of words, to avoid loss of data and to preserve ordinal word position. Thus, we did a random pairing between sentences with the same number of words, after binning the sentences according to their word count. Sentences consisting of 9, 14 or 15 words were infrequent, with fewer than 5 occurrences each. After each permutation, we performed the temporal alignment between sensory modalities (aligning the word onsets), followed by cross-validated MCCA and computation of the time-resolved correlations of crossmodal subject pairs. Due to the long computation time of the canonical variates, we created this null distribution for the exploratory data only. In the figure, color codes for strength of correlation. Colored parcels were most strongly correlated between cross-modal subject pairs (corrected for multiple comparisons).

**Figure 4-1.**
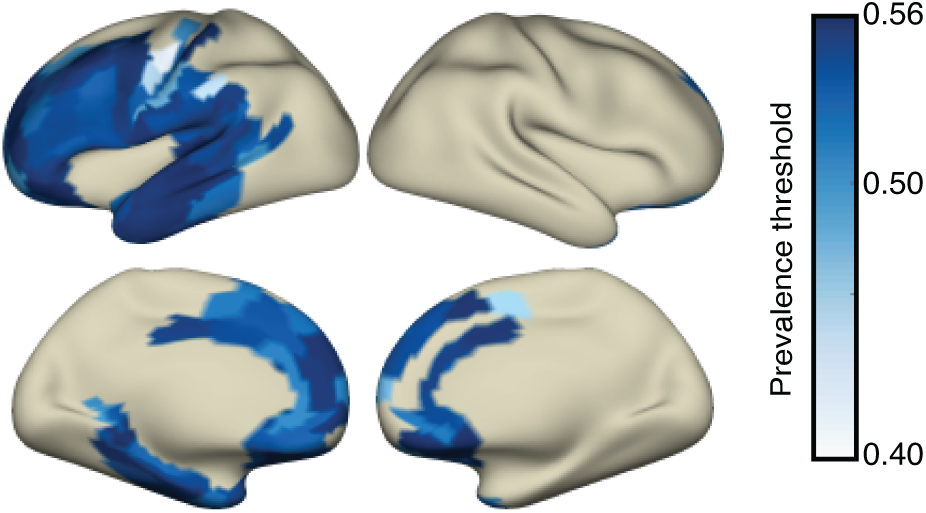
Figure 4-1. Cortical map of maximum threshold *γ*0. For those parcels at which the global null hypothesis could be rejected, the mean (over time) maximum threshold is plotted, for which the null hypothesis can be rejected (*α* = 0.05). Given the sample size of six datasets, the number of second-level permutations and a significance level of *α* = 0.05 the maximally possible threshold that can be reached is 0.5633.

**Figure 4-2.**
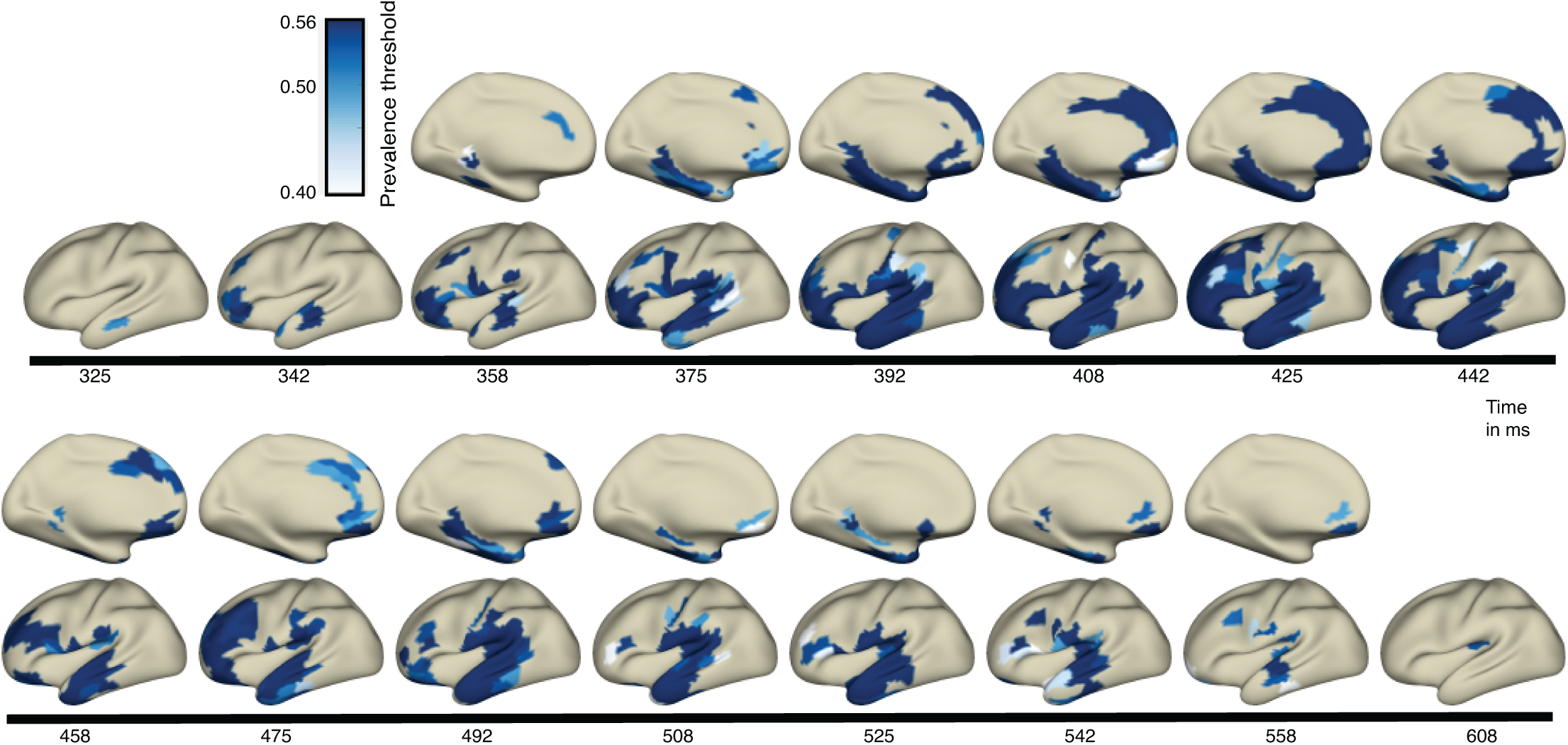
Figure 4-2. Cortical map of prevalence threshold *γ*0. For those parcels at which the global null hypothesis could be rejected, the maximum threshold is plotted, for which the null hypothesis can be rejected at a level of *α* = 0.05.

**Figure 4-3.**
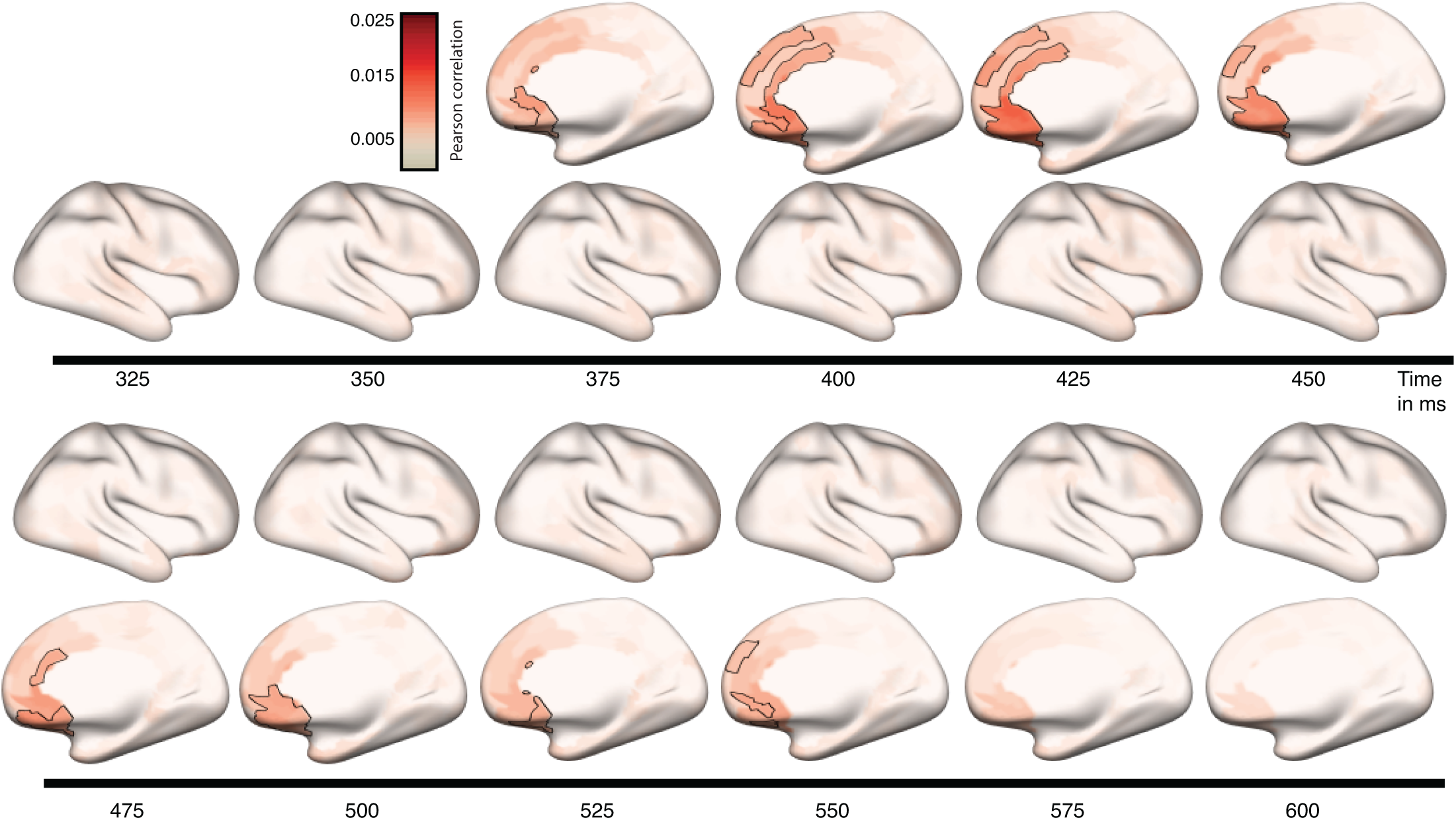
Figure 4-3. Time-resolved spatial maps of cross-modal correlations for the right hemisphere. The average correlation over all six datasets is shown. Color codes for strength of correlation. In addition, the parcels at which the majority null hypothesis according to prevalence inference could be rejected are outlined in black.

